# MpoxRadar: a worldwide Mpox genomic surveillance dashboard

**DOI:** 10.1101/2023.02.03.526935

**Authors:** Ferdous Nasri, Kunaphas Kongkitimanon, Alice Wittig, Jorge Sánchez Cortés, Annika Brinkmann, Andreas Nitsche, Anna-Juliane Schmachtenberg, Bernhard Y. Renard, Stephan Fuchs

## Abstract

Monkeypox (Mpox) is mutating at an exceptional rate for a DNA virus and its global spread is concerning, making genomic surveillance a necessity. With MpoxRadar, we provide an interactive dashboard to track virus variants on mutation level worldwide. MpoxRadar allows users to select among different genomes as reference for comparison. The occurrence of mutation profiles based on the selected reference is indicated on an interactive world map that shows the respective geographic sampling site in customizable time ranges to easily follow the frequency or trend of defined mutations. Furthermore, the user can filter for specific mutations, genes, countries, genome types, and sequencing protocols and download the filtered data directly from MpoxRadar. On the server, we automatically download all Mpox genomes and metadata from the National Center for Biotechnology Information (NCBI) on a daily basis, align them with the different reference genomes, generate mutation profiles, which are stored and linked to the available metainformation in a database. This makes MpoxRadar a practical tool for the genomic survaillance of Mpox, supporting users with limited computational resources. MpoxRadar is open-source and freely accessible at https://MpoxRadar.net.

**GRAPHICAL ABSTRACT:** 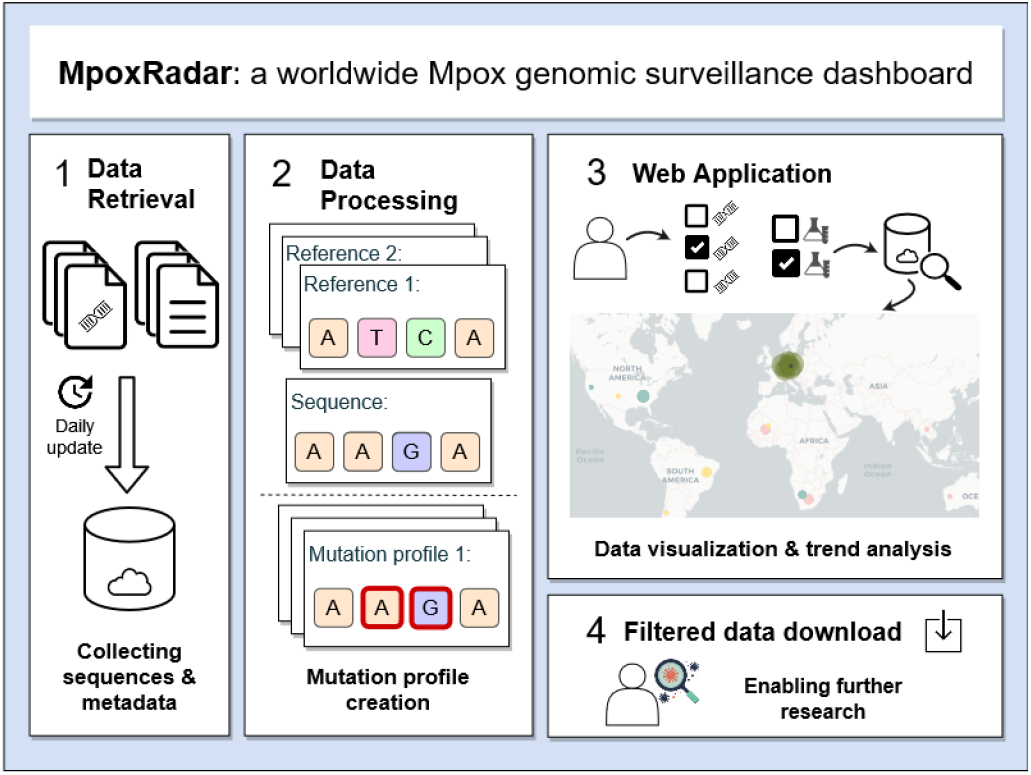

## INTRODUCTION

Mpox (formerly known as Monkeypox) is an infectious disease, with the first recorded infection more than five decades ago in the Democratic Republic of Congo (1). Belonging to the Orthopoxvirus genus, this DNA virus can infect many species. In 2022, Mpox has been declared a public health emergency given its outbreak in non-endemic countries by the World Health Organisation (WHO) with over 86,000 confirmed cases in 110 countries (Global WHO report as of March 10th, 2023).

The SARS-CoV-2 pandemic showed the importance of genomic surveillance in tracking of evolutionary changes to the virus and its spread (2, 3). This also holds for Mpox as Mpox mutated distinctly and changed hosts, creating different clades that caused different spread patterns (4).

Here, we introduce MpoxRadar, a tool for the genomic surveillance of Mpox. It facilitates the interaction and download of genomic sequence and associated metadata through customizable filters, sophisticated queries and intuitive visualisation of results. MpoxRadar was developed with several key analyses in mind that other available systems cannot perform, but that are crucial for end users. To the best of our knowledge, MpoxRadar is the only tool up to date enabling the comparison of sequencing technologies, since this information requires more preprocessing of NCBI metadata and is difficult to obtain. MpoxRadar can assist public health authorities to quickly follow-up on virus variants, even single mutations, or to identify possible sequence technology specific errors such as amplicon dropouts or problems in homopolymer regions. For a comprehensive overview see Tables 1 and 2.

**Table 1.** List of known websites showing Mpox genomic data.

| Name | Description | Website link |
| --- | --- | --- |
| WHO | A long report focused on showing the cases, deaths globally and in different countries. | worldhealthorg.shinyapps.io/mpx_global/ |
| Centers for Disease Control and Prevention (CDC) | Few USA-focused sites with slightly different focuses, i.e., cases, deaths, demographics (gender, race), symptoms, and vaccines. There is one global map with case and death counts. | cdc.gov/poxvirus/monkeypox/response/2022/index.html |
| GISAID | The submission tracker offers a map with a count of available sequences, case count per country, and lineage information. The Variant tracker shows lineage relative frequencies over time along with the count of sequences submitted over time. Further variant tracking site, shows only for 'Clade IIb' the global tracked variant counts (based on submission date), the relative variant genome frequency per continent, and tables for sequence submission with country counts and the name for the most recent sample. There is no link or connection between these sites. | gisaid.org/submission-tracker-global-hmpxv and gisaid.org/hmpxv-variants-dashboard and gisaid.org/hmpxv-variants |
| Nextstrain (5) | Dashboard with a global map showing the present lineages in each country, including a phylogeny tree, with various filters available | nextstrain.org/monkeypox/hmpxv1 and nextstrain.org/monkeypox/mpxv |
| mpoxSpectrum (6) | Dashboard showing number of sequences over time, overall sequence counts per country, overall lineage counts with filters available for country, clade, lineage, host and time. | mpox.genspectrum.org/explore |
| MPoxVR (7) | Global count of Mpox sequences uploaded and country of their origin. There is a sequence search option with country, host, collection and release date, sequence length and completeness filters with tabulated display of results. A variation search is also available with gene names, annotation and mutation type and genome position filters. For this filter there are visualisation options, however they failed to load for all our searches. Lack of a help page makes usage or understanding of filters a sizable challenge. | ngdc.cncb.ac.cn/gwh-poxvirus/ |

**Table 2.**
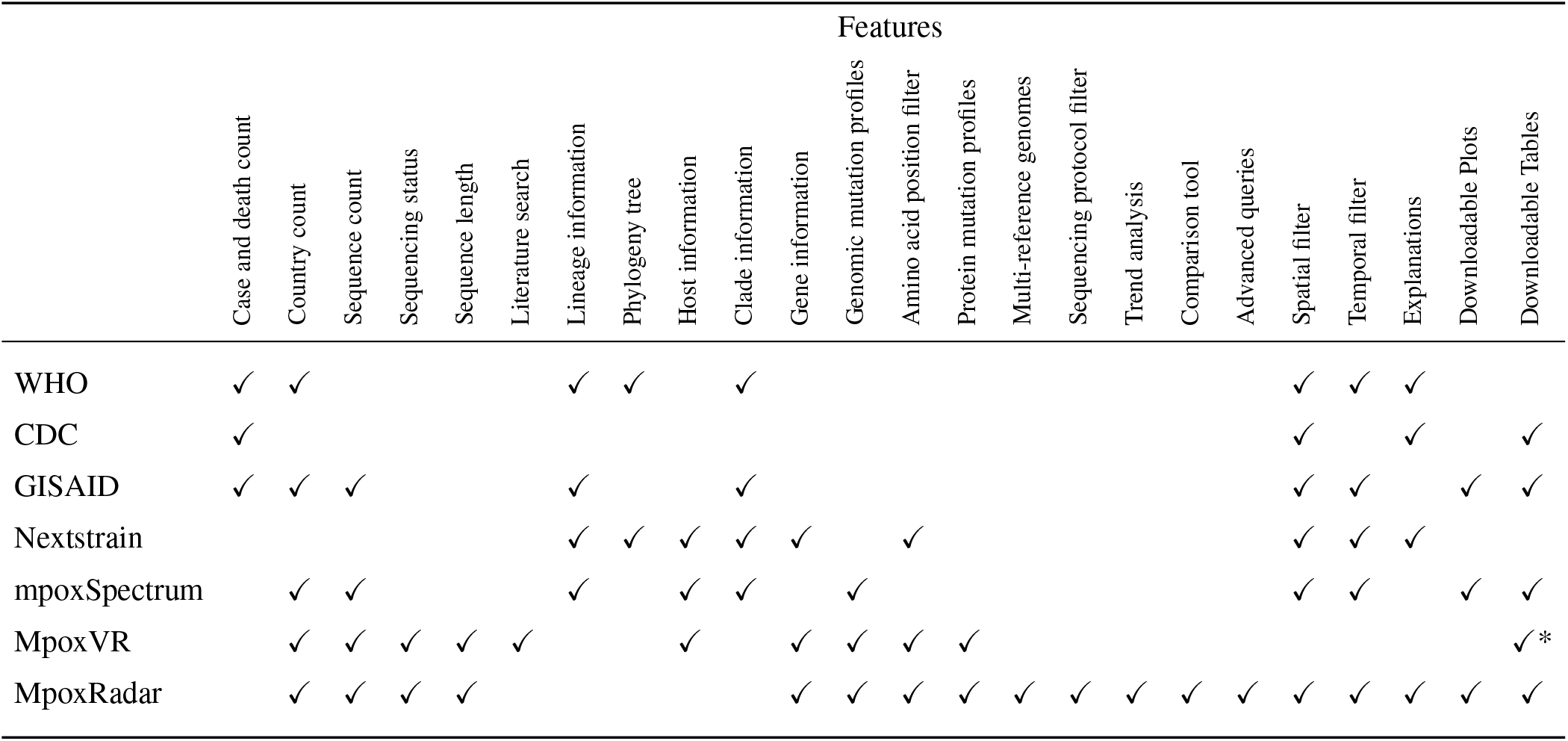
Selected features of websites mentioned in Table 1 and our tool, MpoxRadar, compared to one another. Here, we looked at only the global and interactive sites (when more than one was available). Note, GISAID’s feature list is an accumulation of features across three different sites, which unlike other tools, e.g. MpoxVR or MpoxRadar, are not interlinked. * only limited download size enabled.

## MATERIALS AND METHODS

### Overview

As shown in Figure 1, our web server is supported by an efficient back end system. To ensure security, process, and resource independency, we use three separate Linux servers on the de.NBI cloud infrastructure (cloud.denbi.de).

**Figure 1.**
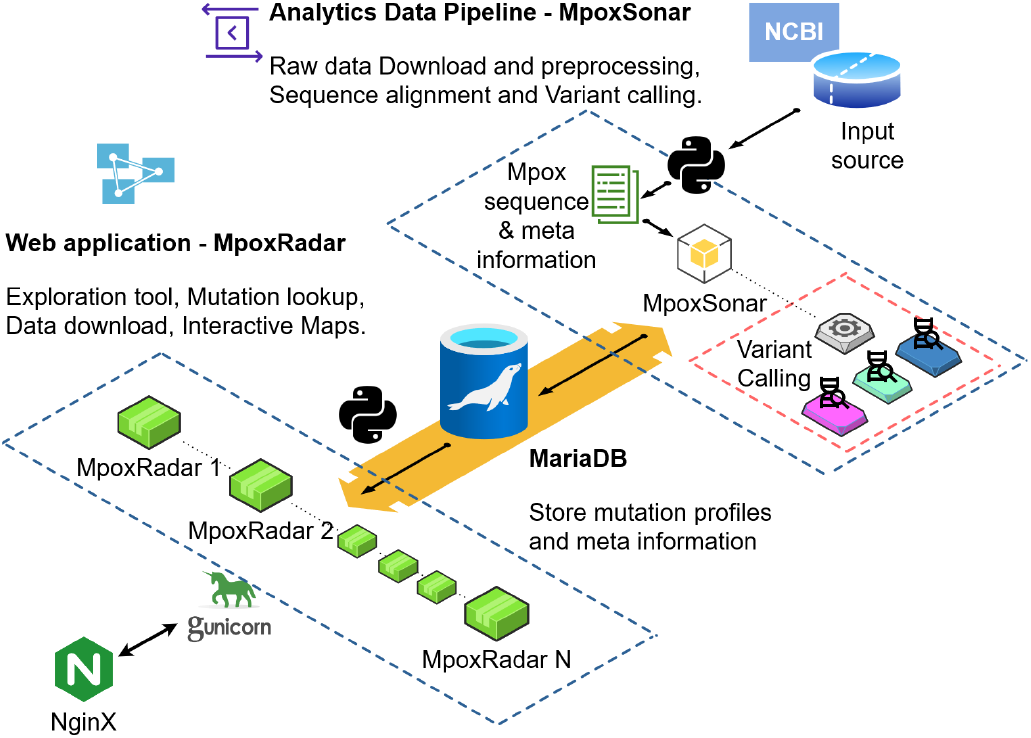
Simplified overview of the processes and steps taken to obtain, process, store and visualise the data on MpoxRadar. The first steps (top right corner) include retrieving data from NCBI, pre-processing, and aligning sequences, followed by variant calling, all done by our analytical tool, MpoxSonar. The data is then stored in a database. Visualisations and interactive maps are presented based on this data on MpoxRadar, while also allowing direct user interaction to the database via the website.

#### Data retrieval

The first server retrieves new samples in Genbank format from the nucleotide database at NCBI on a daily basis (ncbi.nlm.nih.gov/nuccore) using the keyword search `Monkeypox virus[Organism]`. After that, we use the Biopython library (biopython.org) to extract meta information from the files, such as genome completeness, country, and collection date.

#### Pre-processing data

The retrieved data then is cleaned and formatted for downstream processing. This includes the transfer of meta information to harmonized formats. As example, samples are submitted from various countries and labs following different date formats, e.g. using different separators, or orders of date, or even not including a month or day (e.g., *Nov/2022/01* and *09-May-2017*). Thus, all dates are converted to ISO 8601 format (YYYY-MM-DD). If day information is missing, the first day of the given month is used. If no year is given, an empty date is assigned. This along with other cleaning methods help to ensure that the metadata remains consistent and complete for downstream analysis or comparison with other datasets. For import into the database, the DNA sequences are stored in FASTA files and the associated metadata in tab-delimited text files.

#### MpoxSonar

Pre-processed sequences and meta information are further processed by MpoxSonar. The pre-processing and this pipeline run on a second independent server. MpoxSonar is a fork of covSonar, the database-driven system for handling genomic sequences of SARS-CoV-2 and screening genomic profiles (github.com/rki-mf1/covsonar), with additional support for multiple reference genomes and MariaDB as database (mariadb.com). MpoxSonar applies EMBOSS Stretcher (8) to align sequences to defined reference genomes (*optimal global alignment*). Based on the resulting alignment, variations between query and the respective reference sequences are identified (SNPs, deletions, insertions). In the next step, the genetic variations are used to identify protein coding changes. For this, the reference genome annotations are provided as NCBI GenBank files.

Finally, the resulting nucleotide and protein mutation profiles are stored in MariaDB to enable speedy querying. The pipeline has been automated to run daily and processes only new samples to reduce processing time.

#### MpoxRadar

The web application, https://MpoxRadar.net, is written in Python and Dash (github.com/plotly/dash) and communicates with the back end via the database. To allow for optimised usage and data safety, we have the web server running on the third and last independent server. We use Gunicorn (gunicorn.org) as production server together with NGINX (nginx.org) as reverse proxy.

### Multiple Mpox references facilitate mutation analysis

The high mutation rate of Mpox led to multiple clades and distinct lineages over time (9). As a result, more than one reference genome is used for research. We included three different reference genomes in order to facilitate mutation analysis in different locations and points in time. These include genomes with the GenBank Accession numbers NC_063383.1 (identical to MT903340, (10)), MT903344.1 (isolate: MPXV-UK_P2, (10)), and ON563414.3 (isolate: MPXV_USA_2022JMA001, ncbi.nlm.nih.gov/nuccore/ON563414.3). NC_063383.1 is one of the genomes marked as a reference on NCBI. MT903344.1 and ON563414.3 are of interest as they are often used as reference genomes in research (7, 11). When necessary, other references can be easily added by MpoxSonar.

### World map visualises the geographic distribution of mutations

In order to make it easy for users to browse through the data available in our database, our tool page offers numerous filters that can be easily applied by point&click. The user can choose which reference genome they want to use as basis for genomic profiles. According to the selection, the gene filter is updated, the chosen genes then in turn affect an update in the amino acid mutation filter. Furthermore, the user can decide if they want to visualise the frequency of mutations or the increasing trend of their occurence. The trend analysis is based on linear regression using user defined time intervals (default is one month). The time of sample collection, as well as the countries of origin, can also be chosen here.

Moreover, we have a filter for the sequencing technology used to sequence the samples shown, e.g., Illumina, Oxford Nanopore, IonTorrent. This information is crucial as there are innate differences between these sequencing approaches and their biases.

The filters take effect immediately. Resulting frequencies of matching genomes in different geographic locations and time ranges are visualised on the global map, which shows the geographic distribution of the mutation counts. Below the map, there is a bar plot which reacts if the user clicks on a certain country by showing the mutation distribution of that country, at the chosen time and following all the aforementioned filters. Furthermore, the trend or frequency, depending on the user’s choice, of those mutations is plotted separately. All the information shown chosen by the filters is also presented as a table and can be downloaded as a CSV file.

### Compare tool adds comparative analysis functionality

As often in research, it is important to be able to compare the differences between two datasets. In order to simplify this, we created a ‘‘compare” tool that allows users to choose two sets of filtered data sets and get a list of mutations they have in common, and separate lists for mutations that are unique to each dataset. The three sets of data are available in tables for download.

### Database GUI enables advanced and complex queries

MpoxRadar provides the interface to interact with the MpoxSonar query engine and database for specific query types via the ‘advanced” tool tab. For security reasons, we only allow specific MpoxSonar commands listed in Table S1.

Mpox data can be accessed by a keyword or customized search through MpoxSonar. Customized search fields include mutation profile (e.g., “`--profile`”) and properties (always in uppercase, e.g., “`--COUNTRY`”, “`--RELEASE_DATE`” or “`--LENGTH`”). Some examples of commands can be found in Supplementary Table S2. Further customized searches are also accepted, such as “--count” to count all results, or “`--RELEASE_DATE 2022-01-01:2022-01-31`” to select all samples in January. The result of the search will be shown in a table and world map under the advanced tool tab. The interactive map shows the accumulated mutation distribution geographically and can be downloaded. The table of results is also downloadable to enable downstream analyses.

## RESULTS

### Streamlining data gathering chained to MpoxSonar drastically reduces user time in obtaining results

Our database contains all available Mpox nucleotide sequences, complete and partial, downloaded from NCBI. As of March 10th, 2023, this includes 1,862 samples passing our quality filters from across 26 countries. The automatic data download is followed by data cleaning and pre-processing steps, which prepares the data for alignment to multiple reference genomes and the variant annotation. The results along with other functionalities, e.g., trend analysis via linear regression, are available for interaction and download on MpoxRadar.

We not only centralize genomic sequence with associated genotype data and automise the analyses decreasing the time and resources needed for a user to get to the information, but also extract unique metadata like the sequencing technology information. Thereby, we use in-house methods in our pipeline to ensure the quality of the information available. We have tested our web server on focus groups from diverse backgrounds to ensure an intuitive flow and understandable explanations on the help page. Currently, MpoxRadar is actively used for Mpox molecular surveillance by the German national public health institute.

### MpoxRadar allows easy data exploration and extraction, leading to biological findings

It is crucial to understand how viruses interact, survive and replicate in the host cells. A group of DNA editing enzymes from human cells, called the apolipoprotein B mRNA-editing catalytic polypeptide-like 3 (APOBEC3), have been closely studied in research regarding the Human Immunodeficiency Virus (HIV) and later in connection to a vast number of cancer types. They are a part of human innate immunity and are known to edit viral genetics in order to cause erroneous mutations and stop their replication. However, certain mutation signatures (TC > TT and GA > AA) with links to APOBEC3A enzyme have been seen more frequently in Mpox and are a point of interest as they are suspected to have escalated the microevolution and caused the latest outbreak of Mpox (11).

To look into that suspicion, researchers can visualise the geographic evolution or trend of common mutations with the mutation signature “TC > TT”, e.g., “MPXV-UK_P2-076:S30F”, “MPXV-UK_P2-182:S1590F”, “MPXV-UK_P2-157:S111F”, by simply using the filters on the explore page. The results, as shown in Figure 2, indicate an increasing trend and high frequency of some of these mutations over many countries.

**Figure 2.**
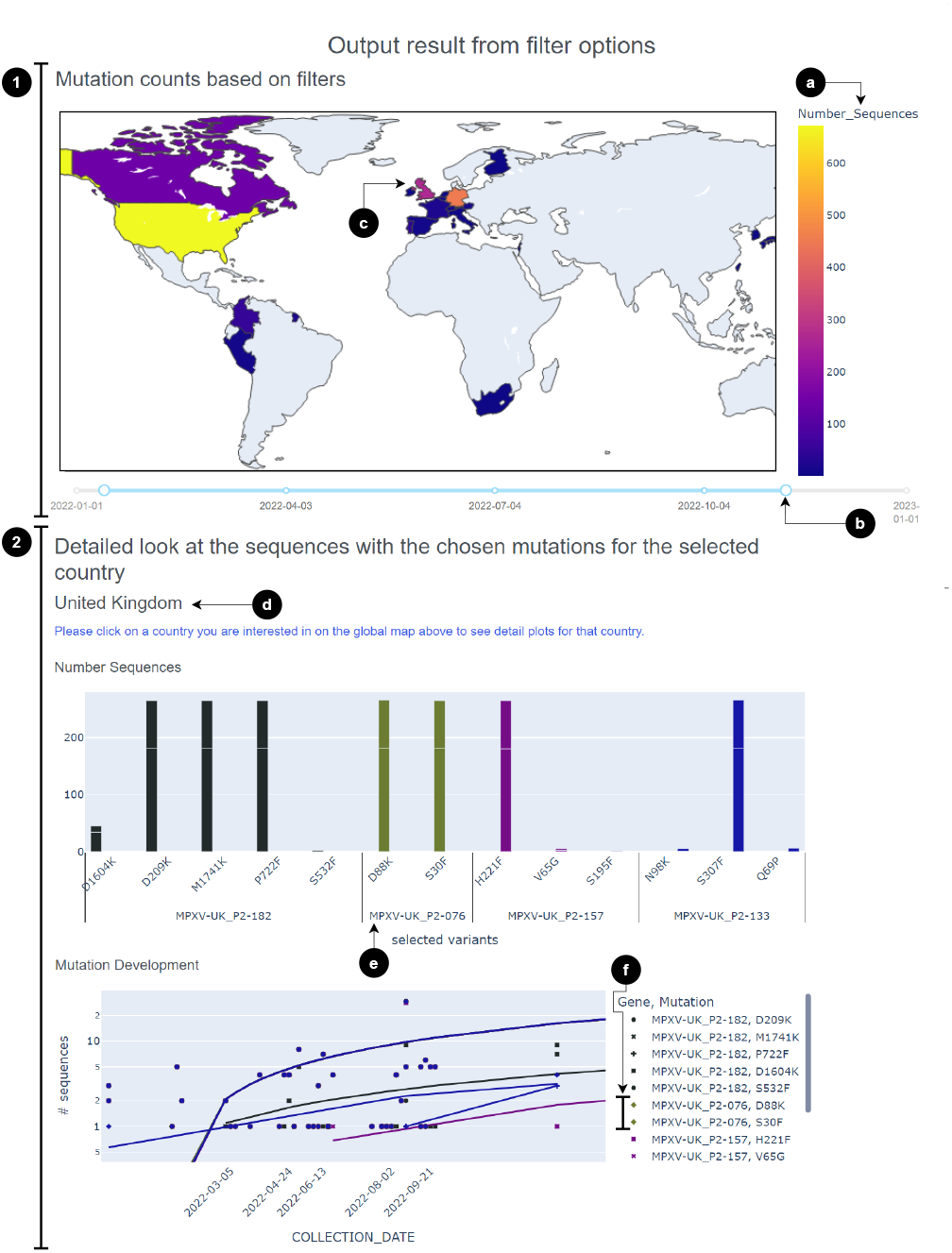
Resulting visualisation from the explore tool, after filtering for the genes “MPXV-UK_P2-076”, “MPXV-UK_P2-133”, “MPXV-UK_P2-157”, “MPXV-UK_P2-182”, and using the pre-selected amino-acid mutation list with the reference genome “MT903344.1”. We changed the interval to 300 days to get a wider coverage of time and evolution of these mutations. The results can be viewed as two sections (marked as 1 and 2). Section 1) The world map with mutation counts provides a quick overview of the presence and frequency of the chosen mutations. (a) The colours on the map represent the number of sequences. One has to keep in mind that some countries sample and publish much more Mpox sequences than other countries, corollary having higher mutation numbers. (b) Users can move the time duration the map is based on by moving this circle. (c) By clicking on a certain country on this map, the users will get detailed information on the mutations in the sequences from that country in section 2 of the output figures. In this case, we click on the United Kingdom, which is also printed to enable a more intuitive understanding of this function for the user at point (d). The first plot shows the number of sequences with chosen mutations on chosen genes. (e) For example, for the gene “MPXV-UK_P2-076”, there are two mutations appearing in the United Kingdom: “D88K” and “S30F” with equally high frequencies. On the mutation development plot below, the user can see the trend of the mutation frequency over time. To increase visual intuition, we colour code each gene with a certain colour starting from the filter input for the gene and its mutations to the plots for sequences including that gene and the mutation development. (f) Here, you can see the colour green used for the gene “MPXV-UK_P2-076” which shows a steady increase in the two mutation frequencies on the plot.

### Comparative analysis between mutations found solely with a certain sequencing technology

Comparative analysis can be simply carried out to find unique mutations given certain filters. For example, if users want to find unique mutations only detected in samples from certain sequencing machines.

As seen with SARS-CoV-2 samples (12), there are biases with each sequencing technology and it is important to allow researchers to quickly and accurately asses differences and to link them to technological and epidemiological information. Using our comparison tool for such a search yields, i.a., 12 counts of the unique Oxford Nanopore mutation “A196593AAAA”. This is of interest as it shows an insertion in a homopolymerous DNA region. These are known to cause difficulties in Oxford Nanopore and Illumina Technologies in general and are interesting in Mpox as well (13). To get a deeper look into this, we can use the “advanced” tool to find other similar mutations in this region. As a result, we find there are a few Illumina Adenine insertion mutations at this position as well, although they report lower numbers of inserted ‘A’ nucleotides.

### Complex searches highlighted potential technical artifacts

Using the advanced query tool, users can have more sophisticated searches as shown in the previous section. Another example is to search for uncovered genome consensus termini which are falsely annotated as nucleotide mutation deletions, while they are commonly a result of amplicon primer schemes. These are used to enrich specific DNA regions and must be adapted to the pathogen and to its genetic changes (14). With the command “`match -r NC_063383.1 --SEQ_TECH ‘Oxford Nanopore Technologies’ --profile del:=1-=200 del:=197010-=197209`”, we can find all samples which were sequenced using “Oxford Nanopore Technologies” and have nucleotide deletion mutations in exactly the first and last ‘200’ bp. This search currently finds four sequences, from the same country released on the same date.

## CONCLUSION

Genomic surveillance is a key tool in understanding the evolution and spread of pathogenic outbreaks. Moreover, there is a need for tools which bridge the gap in communication between different expert groups, e.g., biologists, virologists, epidemiologists, and bioinformaticians. We present MpoxRadar, an interactive web service with a powerful underlying infrastructure to allow easy exploration of Mpox sequence data while also not limiting the user in their ability to pose intricate data queries. This tool was built with a strong basis to allow for quick adaptation to sequences from different pathogens, as we enter an era of massive sequence availability and need for quick genomic tracking tools. Furthermore, we want to empower researchers with quick access to downloadable mutation profiles and metadata to understand and help combat disease spread.

## AVAILABILITY

MpoxRadar is a free and open website to all users without a login requirement accessible under https://MpoxRadar.net/. The open-source code is available under github.com/rki-mf1/MpoxRadar for the front end and github.com/rki-mf1/MpoxSonar for the command-line database back end.

## ACKNOWLEDGEMENTS

We gratefully acknowledge support by Ivan Tunov, Injun Park, and Pavlo Konoplev with early stages of our tool development as well as by Jens-Uwe Ulrich for sharing his expertise on Oxford Nanopore sequencing technologies. We have immense appreciation for NCBI and all data contributors that sequence and upload their data, making research like ours possible.

## SUPPLEMENTARY TABLES

**Table S1.** MpoxSonar commands available via the web application.

| Name | Description |
| --- | --- |
| list-prop | View all properties added to the database. |
| match | Query sequences based on profiles or properties. The match command alone gets all mutation profiles from the database. |

**Table S2.**
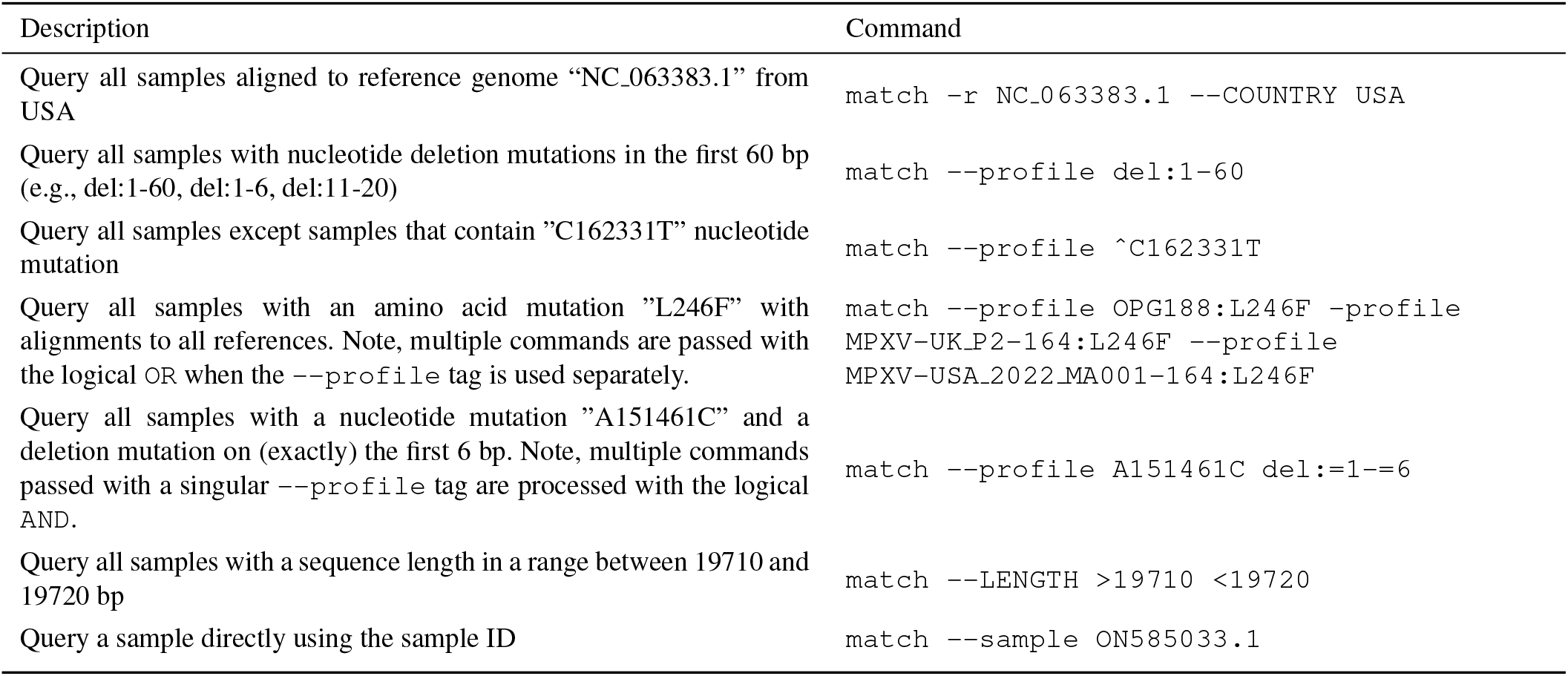
Examples of queries enabled by the MpoxSonar command options.

